# Nanopore sequencing of the pharmacogene CYP2D6 allows simultaneous haplotyping and detection of duplications

**DOI:** 10.1101/576280

**Authors:** Yusmiati Liau, Simran Maggo, Allison L. Miller, John F. Pearson, Martin A. Kennedy, Simone L. Cree

## Abstract

**Background:** The accurate genotyping of *CYP2D6* is hindered by the very polymorphic nature of the gene, high homology with its pseudogene *CYP2D7*, and the occurrence of structural variations. Long read sequencing offers the promise of overcoming some of these challenges, along with the advantage of straightforward variant phasing. We have established methods for sequencing and analysis of DNA amplicons containing the whole *CYP2D6* gene, using the GridION nanopore sequencer.

**Materials and methods:** Seven reference and 25 clinical samples covering various haplotypes including gene duplication were barcoded and sequenced over two sequencing runs. Sequenced raw reads were analyzed using a pipeline of bioinformatics tools including two mapping tools and two variant calling tools.

**Results:** Using minimap2 and nanopolish (mapping and variant calling tools respectively) resulted in the most accurate variant detection. Haplotypes of 52 alleles could be matched accurately to known alleles or subvariants, while the remaining 12 alleles being assigned as novel star (*) allele of novel subvariants of known alleles in the PharmVar *CYP2D6* haplotype database. Allele duplication could be detected by analyzing the allelic balance between the sample haplotypes.

**Conclusion:** Nanopore sequencing of *CYP2D6* offers a high throughput method for genotyping, accurate haplotyping, and detection of new variants and duplicated alleles.

## Introduction

*CYP2D6* is one of the most important and widely studied genes in pharmacogenetics. It codes for the cytochrome P450 2D6 enzyme which is responsible for the metabolism or bioactivation of ∼25% of all available drugs, including antidepressants such as selective serotonin reuptake inhibitors (SSRIs), analgesics such as codeine, and anti-cancer drugs such as tamoxifen [1]. *CYP2D6* is also one of the most polymorphic human genes, leading to considerable variability in enzyme activity between individuals with the need for accurate genotype information for clinical purposes [2].

*CYP2D6* is a relatively small gene (4.4 kilobases from start to stop codon) located on the long arm of chromosome 22. However, the complexities of this gene make accurate genotyping challenging [3-5]. The first challenge lies in the number of variants in the gene. PharmVar (https://www.pharmvar.org/gene/CYP2D6) has compiled more than 100 allelic variations and more than 200 genetic variants [6] in this gene. Moreover, the gnomAD database of more than 140K individuals suggests there may be over 1000 variants in this gene, including more than 400 missense and 30 loss-of-function variants [7].

In addition to its polymorphic sequence, *CYP2D6* can be affected by structural variations including gene deletion, gene duplication, hybrid gene conversion or tandem rearrangement with its upstream pseudogene, *CYP2D7*. Since *CYP2D6* and *CYP2D7* share more than 90% sequence similarity and occur in tandem on the chromosome, care must be taken to selectively target CYP2D6 to obtain accurate genotype information for clinical utility [8, 9].

Various methods have been developed to genotype *CYP2D6*, from the basic restriction fragment length polymorphism to the more sophisticated real time polymerase chain reaction (PCR), microarray, mass spectrometry, or next generation sequencing [10-12]. Most methods are limited by the ability to detect only a subset of pre-selected variants. While next generation sequencing (NGS) methods have the ability to screen all *CYP2D6* variants, these methods are hampered by the risk of misalignment of *CYP2D6* and *CYP2D7* due to the short-reads generated. Most of these methods also do not provide information for variant phasing to confidently determine haplotypes. In practice, this is often carried out using allele specific primers to isolate one or both alleles, followed by Sanger sequencing of the amplification products [13]. This is labour intensive, requires a greater investment of time, and can be technically demanding.

Long read sequencing is envisaged to resolve challenges faced by previous short read technology. For *CYP2D6* genotyping, long amplicon sequencing not only permits unequivocal variant detection without interference from the homologous pseudogene, it also means that variant phasing can be carried out in a reliable and straightforward fashion. A preliminary report described *CYP2D6* sequencing of a single sample using the Oxford Nanopore Technologies (ONT) MinION with an earlier, more error-prone version of the technology [14]. In this paper, we explore the application of current ONT long read sequencing chemistry to simultaneously analyze *CYP2D6* from multiple reference and clinical samples. We performed the sequencing on the GridION platform, a compact nanopore-based instrument from ONT (Oxford, UK) which enables sequencing and data computation in the same module in a real time manner.

## Materials and Methods

### Samples

Thirty-two samples were genotyped over two sequencing runs, with 12 samples in the first and 24 samples in the second run. Four samples from the first run were repeated in the second run as control samples. Seven *CYP2D6* genotyped samples [15] were obtained from Coriell Institute (Camden, NJ, USA) and 25 unrelated clinical samples were collected from two adverse drug reaction cohorts established in our laboratory, Understanding Adverse Drug Reactions Using Genomic Sequencing (UDRUGS) and Genetics of Antidepressants study (GO-A) [16]. These studies received ethical approval respectively from Southern Health and Disability Ethics Committee (URA/11/11/065) and Northern B Health and Disability Ethics Committee (18/NT/21), Ministry of Health, New Zealand. The clinical samples consisted of 14 blood samples and 11 saliva samples. DNA from blood samples was extracted using a modified salting-out method [17] followed by a phenol-chloroform purification step (Supplementary File 1). Saliva samples were collected via an Oragene DNA (OG-500) kit (DNA Genotek Inc., ON, Canada) and DNA was extracted according to the manufacturer’s protocol.

### CYP2D6 Long PCR

The 6.6 kilobases (kb) long PCR amplifying the whole *CYP2D6* gene, including upstream and downstream non-coding regions, was performed according to Gaedigk, 2007 [18]. Specific sequences from ONT were added to the 5’ end of forward and reverse primers to enable sample barcoding in subsequent steps. 250 ng of genomic DNA was amplified in the presence of 1x Kapa LongRange Buffer, 0.3 mM dNTPs, 1.75 mM MgCl_2_, 0.7 μM of each primer, 1 M of betaine, and 1.25 units of Kapa LongRange Taq Polymerase (Sigma-Aldrich) in a 50 μL reaction. PCR was run as follows: 94°C for 3 minutes, 30 cycles of 94°C for 25 seconds, 68°C for 10 seconds, 68°C for 7 minutes, and a final extension at 68°C for 7 minutes.

Deletion and multiplication of the *CYP2D6* gene can be identified by amplifying a 3.5 kb fragment with CYP-13 and CYP-24 primers [19] for deletion and Frag B primers [18] for duplication in a duplex PCR together with the 6.6kb primers. This PCR was carried out as above using 0.4 μM of the 6.6 kb primers and 0.3 μM of the deletion or duplication primers.

In samples where a gene duplication was detected, the duplicated allele was amplified with tailed FragD primers [18] which produced 8.6 kb fragments, and these were subsequently barcoded and sequenced on the GridION. This primer set recognizes the REP-DUP region, a unique combination of *CYP2D6* and *CYP2D7* downstream regions commonly found in duplications of the CYP2D6 gene. This PCR was similar to that for *CYP2D6* described above, except 0.4 μM primers and extension for 10 minutes instead of 7 minutes were used. The sequences of all primers used in this study are listed in Table 5.

### Barcoding PCR

Specific barcodes were assigned to the 6.6 kb PCR products of each sample and 8.6 kb PCR products (where applicable) using the PCR Barcoding Expansion 1-96 kit (EXP-PBC096) (ONT). Barcodes were added using a second-round PCR to 0.5 nM of CYP2D6 6.6 kb or 8.6 kb purified PCR product in the presence of 1x LongAMP PCR buffer, 2 mM of MgCl_2,_ 0.3 mM of dNTPs, and 5 units of LongAmp Taq DNA polymerase in 50 μL reaction. PCR was performed as follows: 95°C for 3 minutes, 15 cycles of 95°C for 15 secs, 62°C for 15 seconds, 65°C for 7 minutes (6.6 kb products) or 10 minutes (8.6 kb products), with a final extension of 65°C for 7 minutes or 10 minutes.

### Modified Beads Purification

Both initial and barcoding PCR products were purified using magnetic beads (HighPrep™ PCR, MagBio, Gaithersburg, MA, USA). In the first sequencing run, we observed that many short (∼1 kb), off-target DNA fragments were sequenced. Therefore, for the second run, to remove any DNA fragments shorter than 3-4 kb, modified beads were made based on the protocol of Ramawater and Schwessinger [20]. In brief, Magbio beads (e.g. 5 mL) were pelleted and re-dissolved in buffer containing 10 mM Tris-HCL, 1 mM EDTA pH 8, 1.6 M NaCl, 11% (w/v) PEG 8000, 0.20% (v/v) Tween-20, and Millipore water to make 5 mL bead solution. For each sample, 0.8x volume of beads was used.

### Nanopore Sequencing Library Preparation

Each DNA library was prepared according to the 1D PCR barcoding (96) amplicons ONT protocol, SQK-LSK109. Briefly, after pooling the barcoded amplicons, 200 fmoles of the DNA pool were subjected to end-repair and ONT adapter ligation. After purification, approximately 350 ng (∼80 fmoles) were introduced to the R9.4 flow cell to be sequenced for 48 hours on the GridION.

### Nanopore Sequencing Data analysis

DNA sequences were extracted from fast5 files using Guppy v2.2.2 with flip-flop basecalling (ONT, Oxford, UK). Only pass reads (qscore > 7) were used for subsequent genotyping analysis as outlined in Figure 1. Guppy basecaller did not accommodate demultiplexing or length filtering when we performed this analysis, and these were done subsequently using Porechop (https://github.com/rrwick/Porechop) and NanoFilt [21]. Filtered fastq files obtained from NanoFilt were then mapped to reference sequence (NG_008376.3, 8593 bps) using minimap2 [22] or NGMLR [23], and variants were called from the mapped Binary Alignment MAP(BAM) files to a Variant Calling Format (VCF) file using nanopolish [24] or Clairvoyante [25]. Threshold for calling variants was set at 0.2 for nanopolish and 0.25 for Clairvoyante as recommended. A trained model for ONT was also used when calling variants using Clairvoyante. Variant phasing in samples containing more than one heterozygote variant was carried out using WhatsHap [26] with maximum coverage of 25X, resulting in phased VCF and phased BAM files. Quality control metrics were obtained using NanoOK [27] and MinIONQC [28].

**Figure 1.**
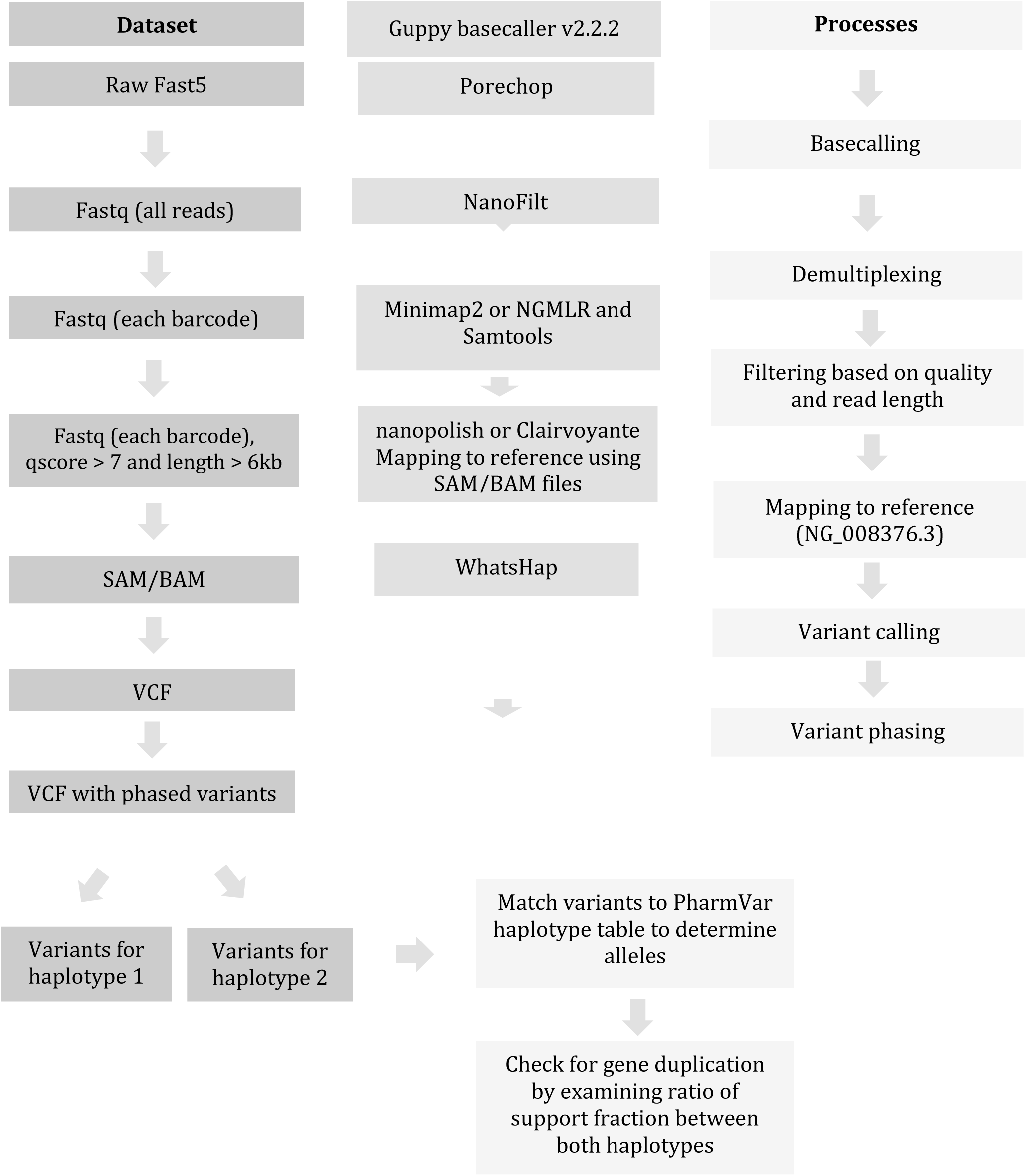
Bioinformatics workflow for *CYP2D6* nanopore sequencing

Variant nomenclature follows the numbering based on NG_008376.3 counting from the sequence start. Haplotype information for each allele was determined by comparing the variants in each allele to the PharmVar haplotype table. Duplicated alleles were identified by examining the ratio of allele frequency between variants from the two alleles of each sample. The presence of duplicated alleles was indicated by a skewed allele balance from 1:1 to 2:1, or more in the case of multiple copies.

Genotype information for reference samples was available through the published Get-RM dataset [15]. Genotype information for the clinical samples was determined using Sanger Sequencing based on key variants (Table 6).

### Sanger sequencing

Nested PCRs covering different regions on the 1:1000 diluted 6.6 kb *CYP2D6* PCR product were performed and used for Sanger sequencing on an AB3130XL Genetic Analyzer (Thermo Fisher Scientific) (Table 5). Nested PCRs were carried out in 20 μL reactions containing 1x PCR buffer, 1.5 mM MgCl_2_, 0.2 mM dNTPs, 0.2 μM of each primer, 0.5 unit of TAQ-TI Heat-Activated DNA Polymerase (Fisher Biotec, Wembley, WA, Australia) and 2 uL of diluted PCR product. DNA was amplified in a touchdown PCR consisting of 15 cycles of 94°C for 15 seconds, 70°C for 15 seconds, and 72°C for 1 minute with annealing temperature decrease of 1°C /cycle followed by 20 cycles of 94°C for 15 seconds, 55°C for 15 seconds, and 72°C for 1 minute and a final extension at 72°C for 5 minutes. Sanger sequencing was performed using 1 μL product of nested PCR, 1 μL of primer, 1x sequencing buffer, and 0.5 μl of Big Dye Terminator (ThermoFisher Scientific, Carlsbad, USA) in 10 μL reaction. Dye incorporation was performed in a thermal cycler with 25 cycles of 96°C for 10 seconds, 50°C for 5 seconds, and 60°C for 4 minutes, and products were subsequently purified using Sephadex G-50 (Sigma-Aldrich, Saint Louis, USA).

## Results

Both sequencing runs resulted in similar yield (∼3.6 gigabases), but different numbers of reads. The first sequencing run resulted in more than 90% reads having length less than 6 kb, possibly due to unspecific amplification during barcode PCR. In the second run, short reads were significantly reduced and target sequences of 6 kb were enriched (∼55% of total reads) with the use of modified beads during PCR product purification (Figure 2). Although the first run resulted in more reads in total, there were more barcoded reads with with >6 kb length in the second run (35% compared to 4%). Despite the differences in number of reads between runs and between samples, all samples had adequate coverage for subsequent analysis and genotyping, with the lowest sample coverage at 235x (Table 1).

**Figure 2.**
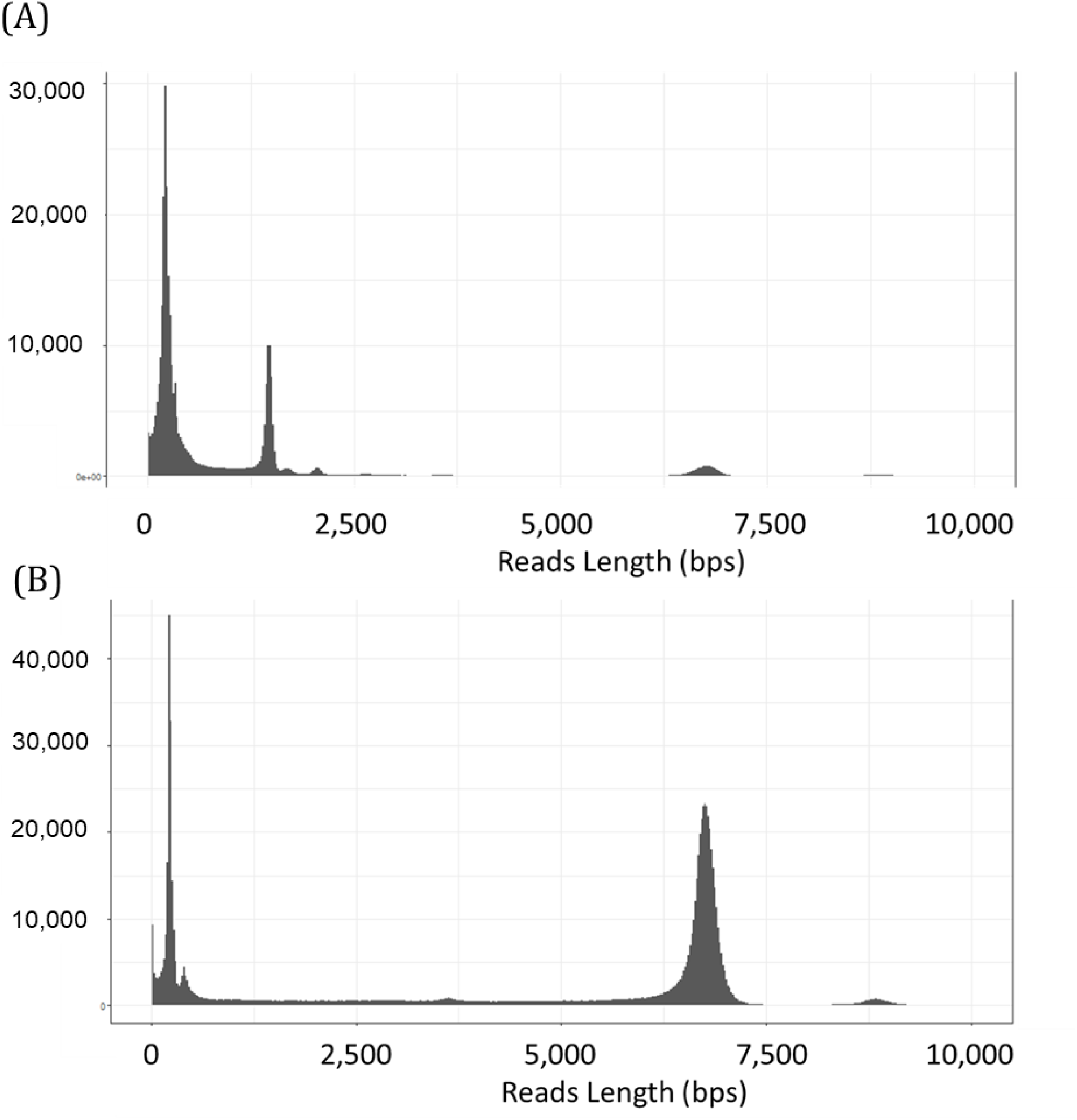
Read Length distribution of (A) the first run, and (B) the second run of *CYP2D6* sequencing

**Table 1.** Metrics of first and second CYP2D6 sequencing runs

| No. | Parameters | 1<sup>st</sup> run | 2<sup>nd</sup> run |
| --- | --- | --- | --- |
| 1. | Total yield | 3.24M reads (3.63 Gbases) | 748K reads (3.62 Gbases) |
| 2. | Read length | N50: 2955 bps<br>Median: 307 bps | N50: 6741 bps<br>Median: 6437 bps |
| 3. | Number of barcodes | 14 | 26 |
| 4. | Number of barcoded reads | 690K reads (47.6% of total pass reads) | 242K reads (46% of total pass reads) |
| 6. | Number of barcoded reads with length > 6kb | 57K reads (3.96% of total pass reads) | 186K reads (35.4% of total pass reads) |
| 7. | Number of reads per barcode (length > 6 kb and qscore > 7) | 235-11832 | 555-21853 |
| 8. | Mean qscore<sup>a</sup> | 6.1 | 8.1 |
| 9. | Error rates (subs – del – ins)<sup>b</sup> | 6.53% - 4.56% - 5.13% | 4.99% - 3.97% - 3.78% |

<sup>a</sup> Mean qscores were obtained from MinION QC
<sup>b</sup> Error rates were obtained from NanoOK

### Comparison of Mapping and Variant Calling Tools

Amplicon data analysis was performed by comparing two mapping tools, minimap2 [22] and NGMLR [23], and two variant calling tools, nanopolish [24] and Clairvoyante [25], resulting in a total of four different analyses. These comparisons were carried out with seven reference samples from the first sequencing run. False positive and negative variants were confirmed by visually inspecting the BAM file on Integrative Genomics Viewer (IGV, Broad Institute, Boston, MA) [29]. The minimum negative predictive value was 99.96% (NGMLR – nanopolish) and the minimum accuracy was 99.68% (NGMLR – Clairvoyante). The calculation for these two measurements was dominated by the relatively high number of true negative variants (6514 – 6561 variants). Clairvoyante generally generated more false positive variants compared with nanopolish, resulting in higher False Discovery Rate values (Table 2). In addition, we observed a misalignment event in NGMLR in the intron 1 CYP2D7 conversion region (seven SNPSs, 4414-4445) (Figure 4). Due to this misalignment, the combination of NGMLR and nanopolish inaccurately called 4423C>G and 4427T>C as 4423insG and 4427delT (Figure 4). Based on our analyses, the combination of minimap2 and nanopolish for mapping and variant calling resulted in the highest sensitivity and positive predictive value and lowest false discovery rate compared to the other tool combinations, and hence was used to analyze the entire sample cohort.

**Table 2.** Comparison between mapping and variant calling tools

| Mapping tool | Variant Calling Tool | TP Variants (median[range]) | FN Variants (median[range]) | FP Variants (median[range]) | PPV (%) | FDR (%) | Sensitivity (%) |
| --- | --- | --- | --- | --- | --- | --- | --- |
| minimap2 | nanopolish | 27 [1-32] | 1 [0-4] | 6 [5-7] | 79.12 | 20.87 | 96.43 |
| minimap2 | Clairvoyante | 23 [1-32] | 2 [0-5] | 17 [10-19] | 57.50 | 42.50 | 92.0 |
| NGMLR | nanopolish | 25 [1-30] | 3 [0-6] | 11 [6-13] | 66.52 | 33.48 | 89.29 |
| NGMLR | Clairvoyante | 26 [1-30] | 2 [0-4] | 19 [12-21] | 54.55 | 45.45 | 92.86 |

Abbreviation: TP: True Positive, FN: False Negative, FP: False Positive, PPV: Positive Predictive Value, FDR: False Discovery Rate. PPV, FDR, and Sensitivity were calculated based on median values.

**Figure 3.**
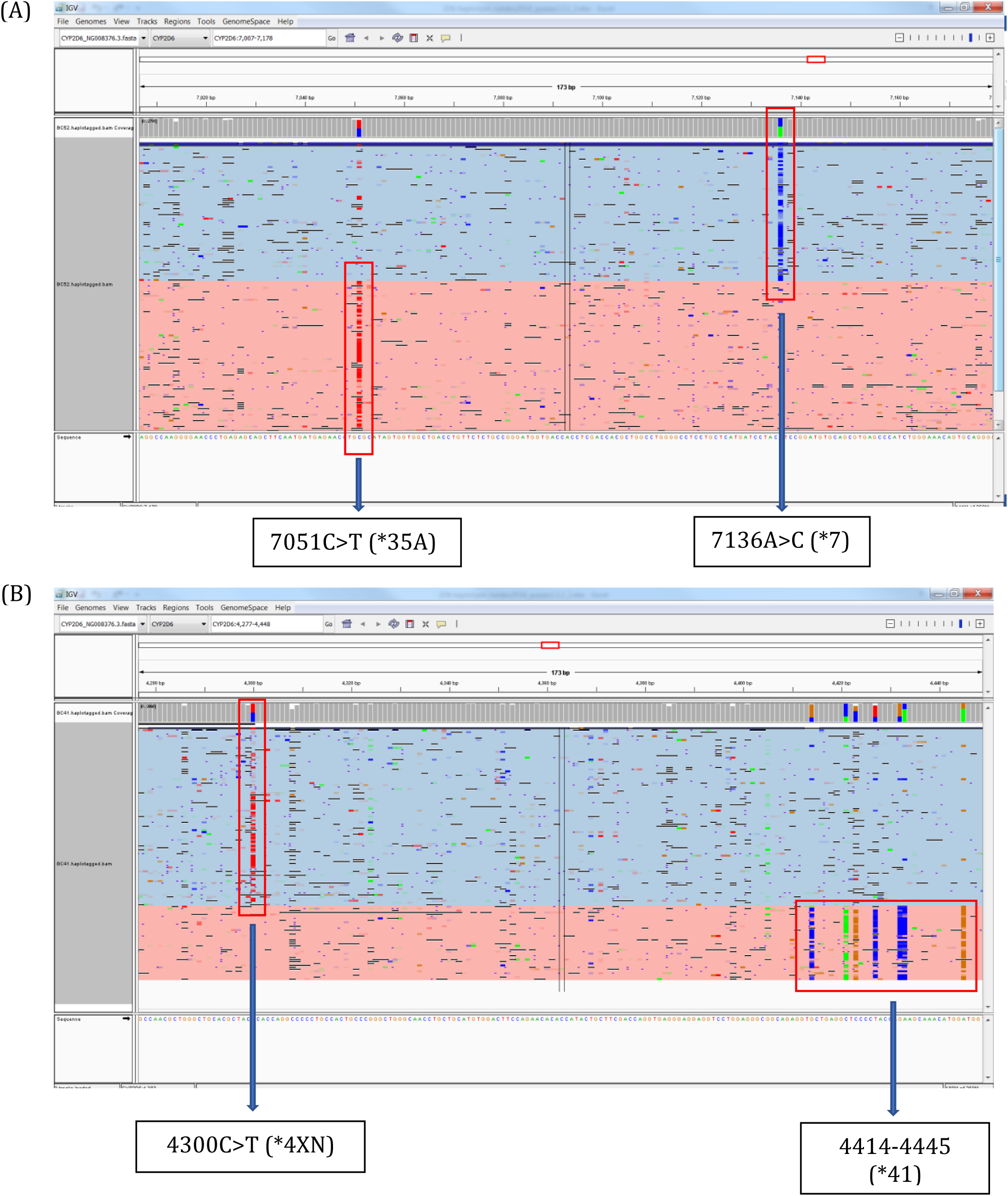
Phased variants for (A) sample with no duplicated allele (NA23348, *7/*35A) and (B) sample with duplicated allele (NA07439, *4XN/*41). Colours represent each of the phased alleles of the samples.

**Figure 4.**
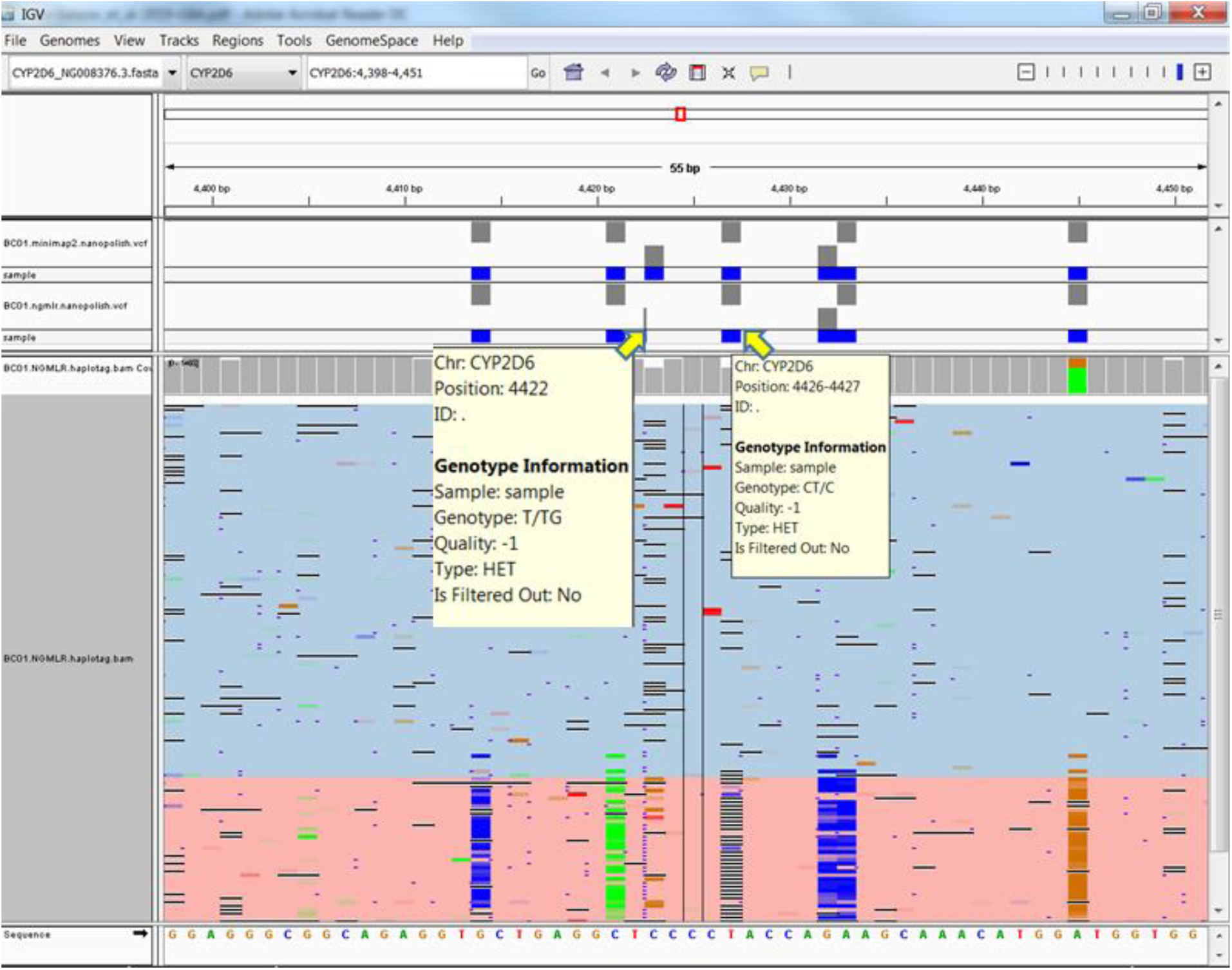
Misdetection of two out of the seven variants of intron 1 CYP2D7 conversion due to misalignment by NGMLR. The two variants were called incorrectly as insertion and deletion (4423insG and 4427delT) instead of the two variants of 4423C>G and 4427T>C.

### CYP2D6 Genotyping Results

Together, the reference and clinical samples cover a total of 16 different star (*) alleles (*1, *2, *3, *4, *6, *7, *9, *10, *15, *17, *28, *29, *33, *35A, *41, and *71), as well as one sample with gene deletion (*5) and four samples with gene duplication. The only sample with a gene deletion (S022) was heterozygous, so only one allele was available to be sequenced. This allele contains a hemizygous variant (6684G>T), which corresponds to *33. This variant was confirmed on Sanger sequencing and the sample was assigned as *5/*33.

The genotyping results for all samples are shown in Table 3. For the 64 alleles represented by the 32 samples, 52 of them were assigned to known subvariant catalogue entries on PharmVar. For the remaining 12 alleles in 11 samples, they either had key variants matched to certain alleles but not to the existing subvariants, or contained variants that did not match any known allele pattern on PharmVar. These, upon submission to PharmVar, were assigned as novel allele or novel subvariants of known alleles. These findings are described below.

**Table 3.** Genotyping Results

| No. | Sample | Sequencing run | Reference Genotype (Get-RM or Sanger) | Allele | Genotype (GridION) | No. of Variants |  |  |
| --- | --- | --- | --- | --- | --- | --- | --- | --- |
|  |  |  |  |  |  | subs | del | ins |
| Reference Samples |  |  |  |  |  |  |  |  |
| 1 | NA07439 | run 1 | *4XN/*41 | hap1 | *4.004XN | 15 | 0 | 0 |
|  |  |  |  | hap2 | *41.003 | 23<sup>a</sup> | 1 | 0 |
|  |  | run 2 | *4XN/*41 | hap1 | *4.004XN | 14<sup>b</sup> | 0 | 0 |
|  |  |  |  | hap2 | *41.003 | 22<sup>a,b</sup> | 1 | 0 |
| 2 | NA23348 | run 1 | *7/*35 | hap1 | *35.001 | 23<sup>b</sup> | 1 | 0 |
|  |  |  |  | hap2 | *7.001 | 1 | 0 | 0 |
|  |  | run 2 | *7/*35 | hap1 | *7.001 | 1 | 0 | 0 |
|  |  |  |  | hap2 | *35.001 | 23<sup>b</sup> | 1 | 0 |
| 3 | NA18518 | run 1 | *17/*29 | hap1 | *17.003<sup>#</sup> | 20<sup>b</sup> | 0 | 0 |
|  |  |  |  | hap2 | *29.001 | 17<sup>c</sup> | 1 | 0 |
| 4 | NA21781 | run 1 | *2/*4(XN) | hap1 | *2.001XN | 22<sup>b</sup> | 1 | 0 |
|  |  |  |  | hap2 | *4.001 | 14<sup>b,d</sup> | 0 | 0 |
| 5 | NA12878 | run 1 | *3/*4 | hap1 | *3.001 | 0 | 1 | 0 |
|  |  |  |  | hap2 | *4.001 | 17<sup>b</sup> | 0 | 0 |
| 6 | NA23405 | run 1 | *1/*7 | hap1 | *7.001 | 1 | 0 | 0 |
|  |  |  |  | hap2 | *1.001 | 0 | 0 | 0 |
| 7 | NA23297 | run 1 | *10(XN) / *17 | hap1 | *17.003<sup>#</sup> | 20<sup>b</sup> | 0 | 0 |
|  |  |  |  | hap2 | *10.002XN | 13<sup>b</sup> | 0 | 0 |
| Clinical Samples |  |  |  |  |  |  |  |  |
| 8 | S001 | run 1 | *1/*1 | hap1 | *1.001 | 0 | 0 | 0 |
|  |  |  |  | hap2 | *1.025<sup>#</sup> | 1 | 0 | 0 |
|  |  | run 2 | *1/*1 | hap 1 | *1.025<sup>#</sup> | 1 | 0 | 0 |
|  |  |  |  | hap 2 | *1.001 | 0 | 0 | 0 |
| 9 | S002 | run 1 | *6/*4 | hap1 | *6.005 | 2 | 1 | 0 |
|  |  |  |  | hap2 | *4.001 | 17<sup>b</sup> | 0 | 0 |
|  |  | run 2 | *6/*4 | hap 1 | *6.005 | 2 | 1 | 0 |
|  |  |  |  | hap 2 | *4.001 | 17<sup>b</sup> | 0 | 0 |
| 10 | S003 | run 1 | *1/*3 | hap1 | *127.001<sup>#</sup> | 9 | 0 | 1 |
|  |  |  |  | hap2 | *3.001 | 0 | 1 | 0 |
| 11 | S004 | run 1 | *4/*9 | hap1 | *4.001 | 17<sup>b</sup> | 0 | 0 |
|  |  |  |  | hap2 | *9.001 | 0 | 1 | 0 |
| 12 | S005 | run 1 | *2/*10 | hap1 | *2.001 | 22<sup>b</sup> | 1 | 0 |
|  |  |  |  | hap2 | *10.002 | 13<sup>b</sup> | 0 | 0 |
| 13 | S006 | run 2 | *2/*9 | hap 1 | *2.001 | 22<sup>b</sup> | 1 | 0 |
|  |  |  |  | hap 2 | *9.001 | 0 | 1 | 0 |
| 14 | S007 | run 2 | *71/*41 | hap 1 | *71.003<sup>#</sup> | 2 | 0 | 0 |
|  |  |  |  | hap 2 | *41.001 | 22<sup>b</sup> | 1 | 0 |
| **15** | S008 | run 2 | *15/*2 | hap 1 | *15.001 | 1 | 0 | 1 |
|  |  |  |  | hap 2 | *2.001 | 22<sup>b</sup> | 1 | 0 |
| **16** | S009 | run 2 | *1/*6 | hap 1 | *6.005 | 2 | 1 | 0 |
|  |  |  |  | hap 2 | *1.025<sup>#</sup> | 1 | 0 | 0 |
| **17** | S010 | run 2 | *4/*10 | hap 1 | *10.002 | 13<sup>b</sup> | 0 | 0 |
|  |  |  |  | hap 2 | *4.015 | 18<sup>b</sup> | 0 | 0 |
| **18** | S011 | run 2 | *2/*2 | hap 1 | *2.001 | 22<sup>b</sup> | 1 | 0 |
|  |  |  |  | hap 2 | *2.001 | 22<sup>b</sup> | 1 | 0 |
| **19** | S012 | run 2 | *4/*4 | hap 1 | *4.001 | 17<sup>b</sup> | 0 | 0 |
|  |  |  |  | hap 2 | *4.001 | 17<sup>b</sup> | 0 | 0 |
| **20** | S013 | run 2 | *2/*41 | hap 1 | *41.004<sup>#</sup> | 23 | 1 | 0 |
|  |  |  |  | hap 2 | *2.001 | 23 | 1 | 0 |
| **21** | S014 | run 2 | *4/*28 | hap 1 | *28.001 | 25 | 1 | 0 |
|  |  |  |  | hap 2 | *4.015 | 19 | 0 | 0 |
| **22** | S015 | run 2 | *4/*35 | hap 1 | *4.001 | 18 | 0 | 0 |
|  |  |  |  | hap 2 | *35.001 | 24 | 1 | 0 |
| **23** | S016 | run 2 | *10/*1 | hap1 | *1.026<sup>#</sup> | 1 | 0 | 0 |
|  |  |  |  | hap2 | *10.003<sup>#</sup> | 14 | 0 | 0 |
| **24** | S017 | run 2 | *35/*41 | hap1 | *41.001 | 22<sup>b</sup> | 1 | 0 |
|  |  |  |  | hap2 | *35.001 | 23<sup>b</sup> | 1 | 0 |
| **25** | S018 | run 2 | *1/*1 | hap1 | *1.016XN | 1<sup>b</sup> | 0 | 0 |
|  |  |  |  | hap2 | *1.012 | 2 | 0 | 0 |
| **26** | S019 | run 2 | *1/*1 | hap1 | *1.012 | 1 | 0 | 0 |
|  |  |  |  | hap2 | *1.001 | 0 | 0 | 0 |
| **27** | S020 | run 2 | *1/*1 | hap1 | *1.025<sup>#</sup> | 1 | 0 | 0 |
|  |  |  |  | hap2 | *1.001 | 0 | 0 | 0 |
| **28** | S021 | run 2 | *1/*1 | hap1 | *1.025<sup>#</sup> | 1 | 0 | 0 |
|  |  |  |  | hap2 | *1.001 | 0 | 0 | 0 |
| **29** | S022 | run 2 | *5/*33 | hap1 | *5.001 |  |  |  |
|  |  |  |  | hap2 | *33.001 | 1 | 0 | 0 |
| **30** | S023 | run 2 | *4/*4 | hap1 | *4.001 | 17<sup>b</sup> | 0 | 0 |
|  |  |  |  | hap2 | *4.026<sup>#</sup> | 18<sup>b</sup> | 0 | 0 |
| **31** | S024 | run 2 | *3/*4 | hap1 | *3.001 | 0 | 1 | 0 |
|  |  |  |  | hap2 | *4.001 | 17<sup>b</sup> | 0 | 0 |
| **32** | S025 | run 2 | *4/*41 | hap1 | *41.001 | 22<sup>b</sup> | 1 | 0 |
|  |  |  |  | hap2 | *4.001 | 16<sup>e</sup> | 0 | 0 |

One sample (S003) had a 6750delA variant in one allele which was called as *3. In the other allele, there are nine SNPs and one insertion variant, which taken together, cannot be matched to any star (*) allele in PharmVar (4470C>T, 7436A>G, 7455T>C, 7466G>A, 7585A>C, 7679A>G, 7685G>A, 7746insT, 7753C>T, 7783A>G). From the 10 variants found in this allele, three of them are in the exonic regions, one of which (7436A>G) is a missense variant, classified as moderate by Variant Effect Predictor (VEP, Ensembl) [30] and is one of the key variants associated with *108. However, the *108 allele has another key variant also in exon 7, 7427A>G (rs61736517), which was not present in our sample. 7466G>A, also a missense variant and predicted to be deleterious by VEP, has been associated with *4.021, but this sample lacks other key variants of *4. This was assigned as a new allele by PharmVar, *127.

One sample (S016) contained only one variant (6504C>T, rs79738337) which is associated only with *60, but lacks the key variant of *60 (6088insTA). As 6504C>T is an intronic variant, this sample was assigned as a new subvariant of *1, *1.026. The other allele of S016 matched *10.002 except for the presence of 4945C>G in this sample. This allele was assigned as *10.003.

Another sample (S007) had two variants (4325G>A and 5695T>C), associated with *71 but lacked the 2617C>G variant and therefore cannot be matched to either *71.001 or *71.002 as per the PharmVar catalogue. This was therefore assigned as a new subvariant, *71.003.

We found five variants not yet catalogued in PharmVar. Two of the Coriell samples (NA18518 and NA23297) with *17 allele had an additional intronic variant (4853C>T, rs376217512). This is a rare variant with allele frequency (AF) of ∼1%. This allele was assigned as *17.003 by PharmVar. One sample with variants matching to *41 allele (S013) contained two non-catalogued variants, 5578C>G (rs143170489) in intron 2 and 5814C>T (rs61736504) in exon 3. Both are rare with AF ∼0.1%. The exonic variant is predicted to be benign when annotated using VEP. This sample was assigned as a new subvariant of *41, *41.004. Another sample with *4 allele (S023) also had a rare (AF: 0.001) intron 2 variant, 5604C>T, rs557722765, and was assigned as a new subvariant, *4.026.

The fifth variant not catalogued in PharmVar was an upstream variant (3836G>A, rs1080992) which was found in four samples, and was the only variant in the particular allele of each sample. This rare variant with an AF of less than 1% [7] has been reported in a non CEU population [31]. No functional information is available for this variant. This allele was assigned as *1.025.

### Detection of duplicated alleles

The presence of a duplicated allele in samples was determined by interrogating the ratio of coverage between alleles. Samples without a duplication were expected to have a 1:1 ratio between alleles, while samples with a duplication were expected have a skewed ratio of ∼2:1 (Figure 3). All four samples which indicated gene duplication during the initial PCR screening showed an uneven allele coverage ratio, and the duplicated alleles could be determined from the over-represented allele. In addition, we sequenced the 8-kb fragments in three out of the four samples with duplication. The 8-kb fragments originated only from the duplicated allele, and aligned to the NG_008376.3 reference sequence from position 3860 to 8593. Sequencing of these fragments revealed the same variants as the duplicated alleles identified from the 6 kb fragments and thus confirmed the duplicated allele in all three samples (NA07439, NA23297, S018) (Table 4).

**Table 4.**
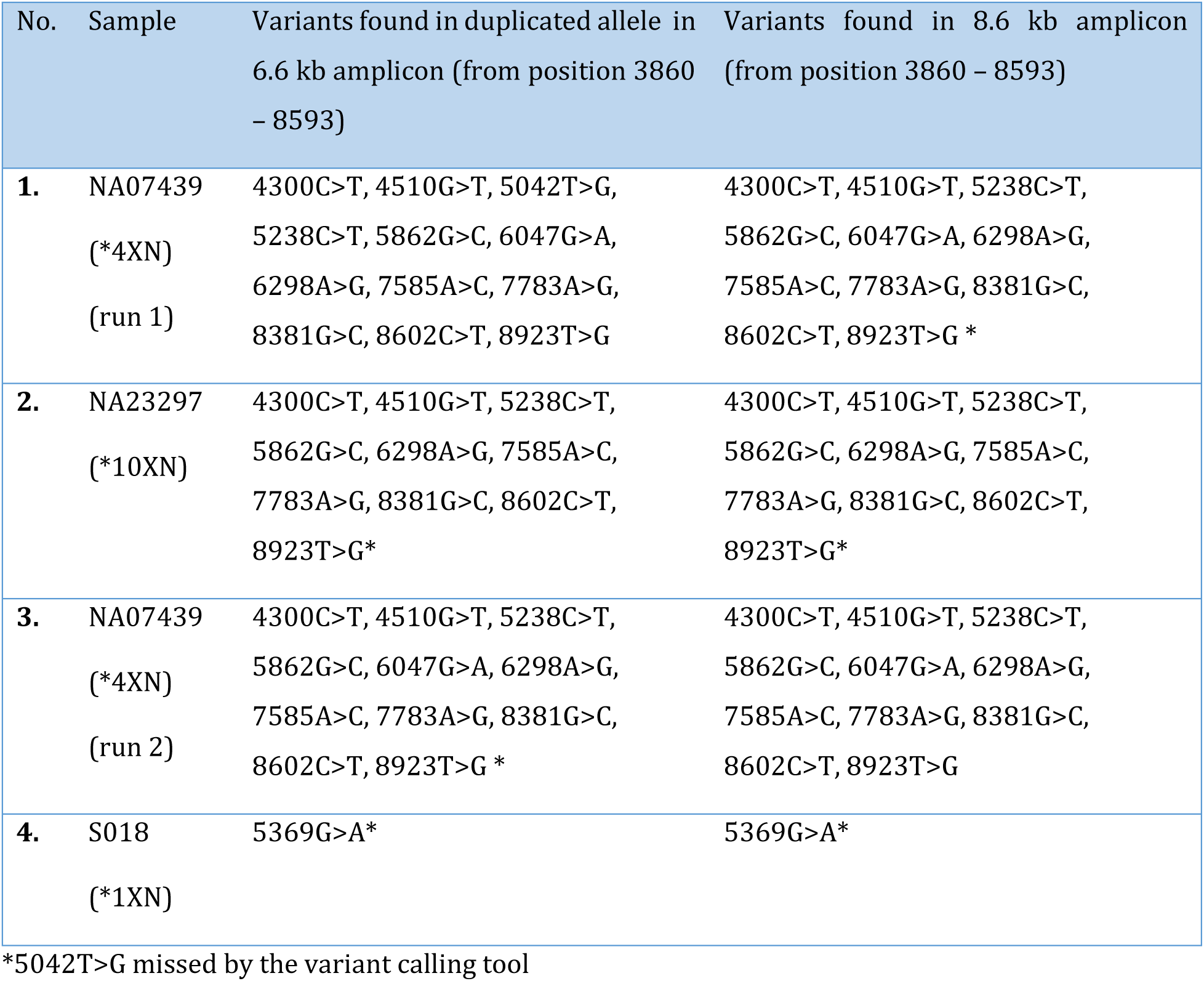
Variants in duplicated allele in 6.6kb and 8kb fragments.

**Table 5.**
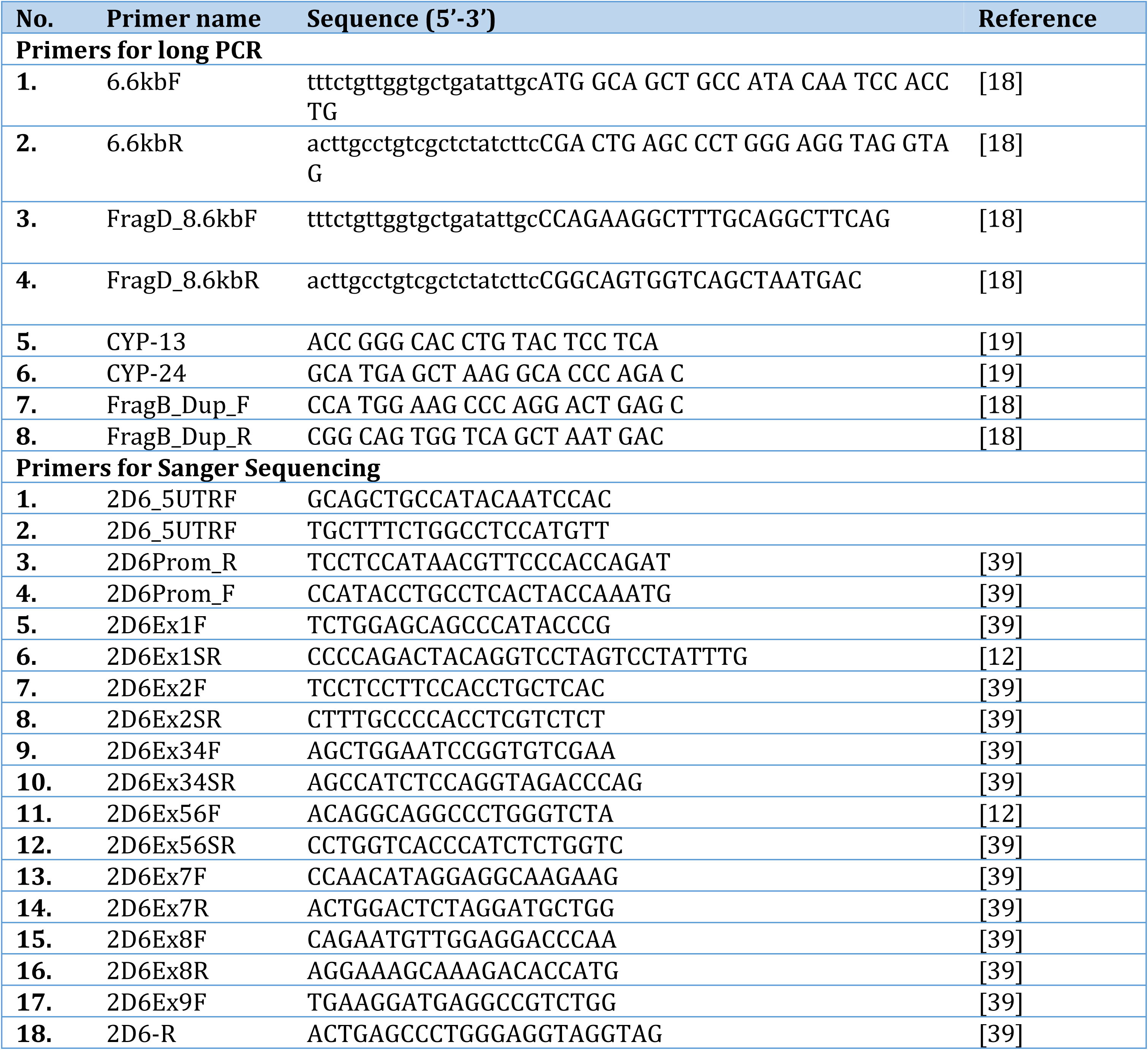
Primer sequences

**Table 6.**
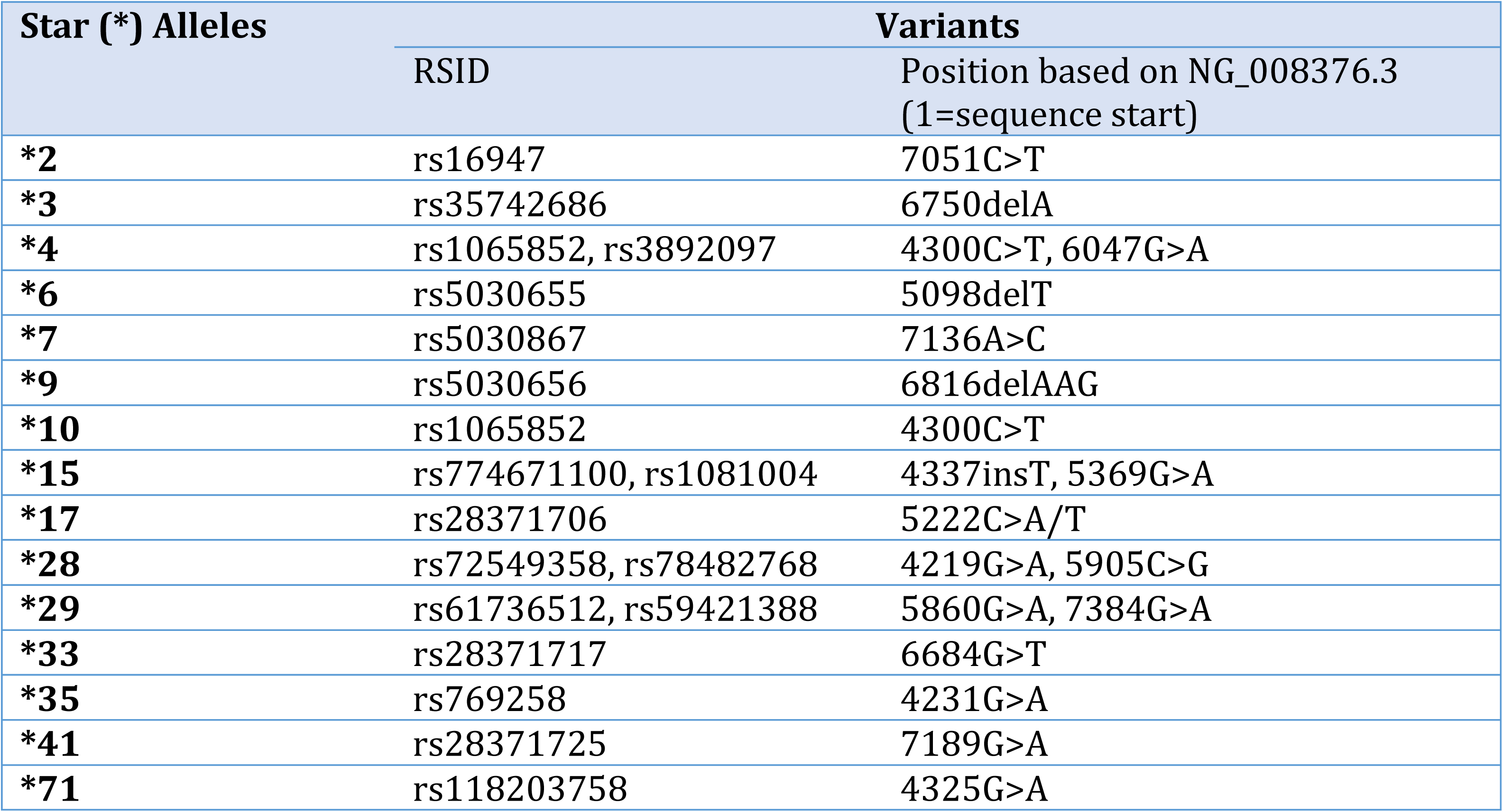
Sequence variants to detect star (*) alleles in Sanger Sequencing

### Validation of duplicated allele and variant phasing using the 8-kb fragment

Interrogating the variants in the 8 kb fragments served two purposes. First, this information identified the duplicated allele as explained in the previous section. Second, comparison of the variants found in the 8-kb amplicon to those found in the 6.6kb amplicon allowed validation of variant phasing performed in the 6.6 kb amplicon. All variants found in the 8-kb amplicons of the three samples matched completely those found on the over-represented (duplicated) allele on the 6.6 kb fragments, thus validated the accuracy of variant phasing of the 6.6kb amplicons. In addition, all phased variants in all samples could be matched to a particular allele or subvariant, confirming the accuracy of variant phasing.

### Identification and Filtering of False Positive Variants

Nanopore sequencing still fails to accurately basecall homopolymer strings. When aligned to the reference, this usually leads to a drop in coverage and variant calling tools report this as a deletion. Two to 13 false positive deletion/insertion events were found in each sample, and all were in regions with three or more sequential identical bases. In addition to errors due to homopolymers, false positive transition variants also occurred at two or three sites per sample. These false positive variants were non-random, always transitions (A↔G or C↔T) and were called only in specific positions: 2381A>G, 2446T>C, 5207C>T, 6440T>C, 8046C>T, 8047C>T, or 8723G>A. These false positive variants have lower quality in the nanopolish produced VCF files compared to the true variants. We applied a previously reported strategy [32] by using quality/coverage ratio to differentiate true and false variants. Most of the true variants have quality/coverage ratio of >1, while most of the false variants have ratio < 0.5. For variants with ratio value between 0.5-1, the trueness of the variant was determined visually in the BAM files using IGV.

Although all samples could be genotyped unequivocally, several variants failed to be detected and were not reported in the VCF file (Table 3). One variant that was present but not reported in many samples was 5042T>G. This SNP sits between four Gs in the upstream and another G downstream, and when the alternate G exists, it created a string of six G’s and the read depth dropped significantly at that site, presumably due to the inability of nanopore sequencing to detect homopolymers correctly. As a result, this SNP was often called incorrectly as a T-deletion instead of a SNP. However, this variant is quite common and is shared by many alleles including the wildtype *1, *2, *4, *10, *41 and many others, and the absence did not interfere with correct haplotyping of any sample.

## Discussion

We performed long amplicon nanopore sequencing of the whole *CYP2D6* gene (including upstream and downstream regions) on the GridION nanopore sequencing platform. To our knowledge, this is the first extensive analysis of *CYP2D6* on a nanopore sequencing platform across multiple star (*) alleles, including assessment of structural variation. An earlier report of *CYP2D6* nanopore sequencing only used one reference sample and was done on the earlier version of ONT chemistry and flow cell. High error rates were associated with that version, and unresolved ambiguity was reported in the sample diplotype, although variant phasing was performed accurately [14]. The initial relatively high error rate associated with nanopore sequencing has improved as the technology continues to evolve. This is mainly attributed to improvements in the pore, sequencing chemistries and bioinformatic algorithms available for analysis. The accuracy of ONT nanopore sequencing has improved from less than 80% with the R7.3 flow cell [33] used by Ammar et. al, to more than 90% with the current R9.4 version [34].

We used the R9.4 flow cell with 1D sequencing chemistry, and generated DNA sequences using the guppy basecaller from ONT. Two different mapping and variant calling tools were then used to analyze the reference samples to select the best bioinformatic workflow for our dataset. We observed that a combination of minimap2 as a mapping tool and nanopolish as a variant calling tool generated the least number of false positive and false negative variants compared to other workflows. The comparison results between nanopolish and Clairvoyante in this study reflected the findings in another study [25], where Clairvoyante reported more false positive variants, and less false negative variants. However, in our study, although Clairvoyante reported some variants missed by nanopolish, it still failed to call several other variants, especially those in the intron 1 CYP2D7 conversion region, resulting in comparable false negative variants between the two tools. Moreover, the quality scores in nanopolish-generated VCF files were more indicative of the trueness of the variants, and false positive variants were more easily filtered.

The advantages of long amplicon sequencing of *CYP2D6* have been recently reported by two papers, using the PacBio single molecule real-time (SMRT) sequencing platform [35, 36]. Although both have the advantages of long read sequencing, the ONT nanopore sequencing platform offers the advantage of very low capital costs and presents a unique opportunity to deliver results in real-time settings such as in the field [24, 37] or at point of care.

One of the issues reported with nanopore sequencing is the limitation in reliable detection of small insertion and deletion (indel) events [32, 38], and we encountered several false positive indels in homopolymer regions. However, true indels outside the homopolymer regions could be detected accurately, including key indel variants associated with common *CYP2D6* star (*) alleles such as 6750delA (*3), 5098delT (*6), 6816delAAG (*9), and 4337insT (*15). All samples with these haplotypes could be identified accurately. In addition, other indel variants were also detected including 4933delC, 7746insA, and 8857delACA which are associated with several star (*) alleles including *2, *41, and *35.

The challenge that homopolymer regions present to the current nanopore sequencing also means that this technology remains limited for investigating the two long homoplymer strings of A’s in the upstream region of the *CYP2D6* gene (2943-2964 and 3098-3107). The regions in between these two strings are notoriously difficult to sequence, resulting in a gap in most PharmVar catalogued alleles or subvariants. Variation in the number of A’s within 2943-2964 has been reported previously [36], however this region has not been included as part of the haplotype definition [6]. Although nanopore sequencing was not able to reliably detect variation in the poly A string, the regions adjacent to the strings were sequenced accurately in this study, including the detection of variants such as 2966A>G, which are still ambiguous in many star (*) alleles.

Sample barcoding permits efficient use of the flow cell and reduction in per sample assay costs. For high throughput laboratories or for research studies on larger cohorts, sample number could potentially be multiplexed up to 96 using the ONT barcode kit. Our 24-plex sequencing run resulted in a minimum read depth of ∼500x, which is more than adequate for accurate variant calling and phasing, while some samples had tens of thousands of reads. The lowest read depth in our sample cohort was ∼200x, and we observed no difference in accuracy compared to samples with higher read depth. Using nanopore sequencing for a long amplicon from the *GBA* gene, Leija-Salazar et al reported that coverage >100x is sufficient for accurate variant calling [32].

The main advantage of long read sequencing is enabling straightforward variant phasing. In this study, all variants could be phased accurately (provided more than one heterozygote variant is present), resulting in highly confident haplotyping to the subvariant level. In addition, duplicated alleles when present, were also identified. Using the 8-kb PCR amplicon derived from duplicated alleles as a confirmation sequence, we showed that duplicated alleles could be detected from the 6kb *CYP2D6* amplicon by the skewed allelic balance of reads. The allele ratio could be observed from the VCF file or by visualizing the phased BAM file on IGV. All our duplication samples have an allele ratio of ∼2:1, indicating two copies of the duplicated gene. Due to limitation of the samples used in this cohort, it remains inconclusive whether this method could determine the exact number of gene copies accurately in cases with more than two copies.

Other challenges remain. The whole gene deletion allele (*5) still requires detection by a separate duplex PCR, and despite the accurate determination of haplotypes and duplication, the methods described here have not been extended to the detection of other CYP2D6 structural variations such as the CYP2D6-CYP2D7 hybrids or tandem rearrangements.

## Conclusion

In summary, nanopore sequencing of the *CYP2D6* gene allows accurate detection and phasing of variants and determination of sample haplotypes to the subvariant level. New variants not yet catalogued on the PharmVar CYP2D6 page (rs1080992, rs376217512, rs143170489, rs61736504, rs557722765) were also reported in several samples, and key indel variants could be reliably detected. In total this study contributed to one novel allele (*127) and seven novel subvariants of known alleles (*1.025, *1.026, *4.026, *10.004, *17.003, *41.004, *71.003). The ability of nanopore sequencing to detect duplicated alleles when present provides better prediction of CYP2D6 phenotype. In addition, sample barcoding permits sequencing and analysis of multiple samples simultaneously, improving cost and time efficiency.

## Acknowlegdements

YL received the University of Otago Doctoral Scholarship. Other funding support was received from Lottery Health Research (New Zealand), Health Research Council of New Zealand, and The Jim and Mary Carney Charitable Trust (Whangarei, New Zealand). We also thank Will Taylor from the Surgery Department, University of Otago, Christchurch for assistance with installation of several bioinformatic tools.

## Supplementary File 1

### Protocol for DNA extraction from blood samples

For each patient sample, 4 mL of peripheral blood was mixed with 6 mL of red blood cell lysis buffer (1 mM EDTA) and spun at 2000 rcf for 10 minutes at room temperature. The supernatant was carefully discarded and the pellet resuspended and the initial step repeated until a layer of white buffy cells is clearly visible. The supernatant was then discarded and 1 mL of 1 M NaCl solution added. This solution was then repeatedly vortexed to eliminate any clumping. Once the solution was clear of clumps, 6 mL of white cell lysis solution (10 mM Tris HCl, 26 mM EDTA, 17.3 mM (0.5%) SDS) and 50 μL of RNase A solution was added and incubated at 37°C for at least 1 hour.

After incubation 1mL of 3M sodium acetate solution was added and mixed. The resulting solution was then added to a 15 mL tube containing 2.5 gr of high vacuum grease (Dow Corning Corporation, USA) and then 1 ml of phenol:chloroform:isoamyl alcohol 25:24:1 was added. The tubes were capped and thoroughly mixed before centrifuging for 5 minutes at 1500 rcf. Five mL of the resultant supernatant was then transferred to a new tube containing 5mL of 100% isopropanol. The tubes were gently inverted until DNA precipitates. The tubes containing DNA were then centrifuged at 2000 rcf for 5 minutes to form a DNA pellet at the bottom of the tube. The isopropanol was carefully discarded and then washed with 10 mL of 70% ethanol (10 mL) and centrifuged at 2000 rcf for 3 minutes. After discarding the ethanol, the DNA pellet was dissolved in 500 mL of TE buffer (10 mM Tris-HCl, 1 mM EDTA, pH 8.0) at 37°C for at least 8 hours. The DNA solution was quantified using a spectrophotometer at 260 nm and diluted to 50 ng/mL.

EDTA: Ethylenediaminetetraacetic acid
TRIS HCL: tris(hydroxymethyl)aminomethane hydrochloric acid
SDS: Sodium dodecyl sulfate
TE buffer: Tris-EDTA buffer
NaCl: Sodium chloride

## Notes

#### Summary of Updates

Novel allele and subvariants have been submitted to PharmVar and assigned a new star (*) allele or subvariant number

## References

1. Ingelman-Sundberg M, Sim SC, Gomez A, Rodriguez-Antona C. Influence of cytochrome P450 polymorphisms on drug therapies: Pharmacogenetic, pharmacoepigenetic and clinical aspects. Pharmacology and Therapeutics 116(3), 496–526 (2007).

2. Gaedigk A, Dinh JC, Jeong H, Prasad B, Leeder JS. Ten years’ experience with the CYP2D6 activity score: A perspective on future investigations to improve clinical predictions for precision therapeutics. Journal of Personalized Medicine 8(2), 1–15 (2018).

3. Gaedigk A. Complexities of CYP2D6 gene analysis and interpretation. International Review of Psychiatry 25(5), 534–553 (2013).

4. Yang Y, Botton MR, Scott ER, Scott SA. Sequencing the CYP2D6 gene: From variant allele discovery to clinical pharmacogenetic testing. Pharmacogenomics 18(7), 673–685 (2017).

5. Nofziger C, Paulmichl M. Accurately genotyping CYP2D6 : not for the faint of heart. Pharmacogenomics 19(13), 999–1002 (2018).

6. Gaedigk A, Ingelman-Sundberg M, Miller NA, Leeder JS, Whirl-Carrillo M, Klein TE. The Pharmacogene Variation (PharmVar) Consortium: Incorporation of the Human Cytochrome P450 (CYP) Allele Nomenclature Database. Clinical Pharmacology and Therapeutics 103(3), 399–401 (2018).

7. Lek M, Karczewski KJ, Minikel EV et al. Analysis of protein-coding genetic variation in 60,706 humans. Nature 536 285–291 (2016).

8. Kimura S, Umeno M, Skodaj RC, Meyert UA, Gonzalez FJ. Sequence and Identification of the Polymorphic CYP2D6 Gene, a Related Gene, and a Pseudogene. Am. J. Hum. Genet. 45 889–904 (1989).

9. Riffel AK, Dehghani M, Hartshorne T et al. CYP2D7 sequence variation interferes with TaqMan CYP2D6*15 and *35 genotyping. Frontiers in Pharmacology 6(JAN), 1-10 (2016).

10. Sachse C, Brockmöller J, Bauer S, Roots I. Cytochrome P450 2D6 variants in a Caucasian population: allele frequencies and phenotypic consequences. American journal of human genetics 60(2), 284–295 (1997).

11. Rebsamen MC, Desmeules J, Daali Y et al. The AmpliChip CYP450 test: Cytochrome P450 2D6 genotype assessment and phenotype prediction. Pharmacogenomics Journal 9(1), 34–41 (2009).

12. Chua EW, Cree SL, Ton KNT et al. Cross-comparison of exome analysis, next-generation sequencing of amplicons, and the iPLEX® ADME PGx panel for pharmacogenomic profiling. Frontiers in Pharmacology 7(JAN), 1 (2016).

13. Gaedigk A, Riffel AK, Leeder JS. CYP2D6 Haplotype Determination Using Long Range Allele-Specific Amplification: Resolution of a Complex Genotype and a Discordant Genotype Involving the CYP2D6*59 Allele. Journal of Molecular Diagnostics 17(6), 740–748 (2015).

14. Ammar R, Paton TA, Torti D, Shlien A, Bader GD. Long read nanopore sequencing for detection of HLA and CYP2D6 variants and haplotypes. F1000Research 4(0), 17 (2015).

15. Pratt VM, Everts RE, Aggarwal P et al. Characterization of 137 Genomic DNA Reference Materials for 28 Pharmacogenetic Genes: A GeT-RM Collaborative Project. Journal of Molecular Diagnostics 18(1), 109–123 (2016).

16. Maggo SDS, Chua EW, Chin P et al. A New Zealand platform to enable genetic investigation of adverse drug reactions. New Zealand Medical Journal 130(1466), 62–69 (2017).

17. Miller SA, Dykes DD, Polesky HF. A simple salting out procedure for extracting DNA from human nucleated cells. Nucleic Acids Research 16(3), 1215–1215 (1988).

18. Gaedigk A, Ndjountché L, Divakaran K et al. Cytochrome P4502D6 (CYP2D6) gene locus heterogeneity: Characterization of gene duplication events. Clinical Pharmacology and Therapeutics 81(2), 242–251 (2007).

19. Steen V, Andreassen O, Daly A et al. Detection of the poor metabolizer-associated CYP2D6(D) gene deletion allele by long-PCR technology. Pharmacogenetics 5(4), 214–223 (1995).

20. Nagar R, Schewessinger B. DNA size selection (>3-4kb) and purification of DNA using an improved homemade SPRI beads solution. doi:dx.doi.org/10.17504/protocols.io.n7hdhj6 (2018).

21. De Coster W, D’hert S, Schultz DT, Cruts M, Van Broeckhoven C. NanoPack: Visualizing and processing long-read sequencing data. Bioinformatics 34(15), 2666–2669 (2018).

22. Li H. Minimap2: pairwise alignment for nucleotide sequences. 34(May), 3094–3100 (2017).

23. Sedlazeck FJ, Rescheneder P, Smolka M et al. Accurate detection of complex structural variations using single-molecule sequencing. Nature Methods 15(6), 461–468 (2018).

24. Quick J, Loman NJ, Duraffour S et al. Real-time, portable genome sequencing for Ebola surveillance. Nature 530(7589), 228–232 (2016).

25. Luo R, Sedlazeck FJ, Lam TW, Schatz MC. A multi-task convolutional deep neural network for variant calling in single molecule sequencing. Nature Communications 10(1), 998 (2019).

26. Martin M, Patterson M, Garg S et al. WhatsHap : fast and accurate read-based phasing. bioRxiv 085050 1–18 (2016).

27. Leggett RM, Heavens D, Caccamo M, Clark MD, Davey RP. NanoOK: Multi-reference alignment analysis of nanopore sequencing data, quality and error profiles. Bioinformatics 32(1), 142–144 (2015).

28. Lanfear R, Schalamun M, Kainer D, Wang W, Schwessinger B. MinIONQC: fast and simple quality control for MinION sequencing data. Bioinformatics doi:10.1093/bioinformatics/bty654(July), 1–3 (2018).

29. Robinson JT, Thorvaldsdoottir H, Winckler W et al. Integrative Genomics Viewer. Nat Biotechnol 29(1), 24–26 (2011).

30. Mclaren W, Gil L, Hunt S et al. The Ensembl Variant Effect Predictor. Genome Biology 17(1), 122 (2016).

31. He X, He N, Ren L et al. Genetic polymorphisms analysis of CYP2D6 in the Uygur population. BMC Genomics 17(1), 1–9 (2016).

32. Leija-Salazar M, Sedlazeck FJ, Mokretar K et al. Detection of GBA missense mutations and other variants using the Oxford Nanopore MinION. bioRxiv doi:10.1101/288068(November 2018), 288068-288068 (2018).

33. Jain M, Fiddes IT, Miga KH, Olsen HE, Paten B, Akeson M. Improved data analysis for the MinION nanopore sequencer. Nature Methods 12(4), 351–356 (2015).

34. Tyler AD, Mataseje L, Urfano CJ et al. Evaluation of Oxford Nanopore’s MinION Sequencing Device for Microbial Whole Genome Sequencing Applications. Scientific Reports 8(1), 1–12 (2018).

35. Qiao W, Yang Y, Sebra R et al. Long-read single-molecule real-time (SMRT) full gene sequencing of cytochrome P450-2D6 (CYP2D6) HHS Public Access. Hum Mutat 37(3), 315–323 (2016).

36. Buermans HPJ, Vossen RHaM, Anvar SY et al. Flexible and Scalable Full-Length CYP2D6 Long Amplicon PacBio Sequencing. Human Mutation 38(3), 310–316 (2017).

37. Quick J, Grubaugh ND, Pullan ST et al. Multiplex PCR method for MinION and Illumina sequencing of Zika and other virus genomes directly from clinical samples. Nature Protocols 12 1261–1276 (2017).

38. Magi A, Semeraro R, Mingrino A, Giusti B, D’aurizio R. Nanopore sequencing data analysis: state of the art, applications and challenges. Briefings in Bioinformatics doi:10.1093/bib/bbx062(June), (2017).

39. Wright GEB, Niehaus DJH, Drögemöller BI, Koen L, Gaedigk A, Warnich L. Elucidation of CYP2D6 genetic diversity in a unique African population: Implications for the future application of pharmacogenetics in the xhosa population. Annals of Human Genetics 74(4), 340–350 (2010).

